# Body Size Variation and Sexual Dimorphism in Eleven Species of Venezuelan Anurans

**DOI:** 10.1101/2024.11.19.624291

**Authors:** Israel Cañizales

## Abstract

This study investigates intra- and interspecific variation in snout–vent length (SVL) and abdominal width (AW) across 11 anuran species in Venezuela, revealing significant patterns of sexual size dimorphism (SSD) in seven species. Females exhibited larger SVL than males in most species (e.g., *Atelopus cruciger*, *Leptodactylus fuscus*, and Phyllomedusa trinitatis), with mean SVL differences ranging from 4 to 15 mm (*p* < 0.05). This study provides the first reference AW values for these species, contributing novel data for morphometric studies. Four species (*Boana punctata*, *Leptodactylus fuscus*, *Rhinella marina*, and *Scinax rostratus*) showed no significant differences in SVL or AW (*p* > 0.05), indicating limited SSD or monomorphism. Statistical analyses highlighted geographic variation in SVL, with female-biased SSD more pronounced in arboreal species and male-biased SSD associated with territorial, burrow-digging behaviors in terrestrial species (*p* < 0.05). These findings support Rensch’s rule, where SSD scales with body size: SSD decreases with increasing female size and increases with male-biased size. In *L. fuscus*, for instance, SVL values (males: 54.5 mm; females: 61.56 mm) exceeded reported ranges but lacked statistical significance (*p* = 0.595), highlighting substantial geographic and ecological influences. Directional asymmetry was evident in arboreal species, linked to their ecological adaptations for locomotion. Conversely, terrestrial species displayed larger size variation associated with competition and reproductive strategies. Statistical tests demonstrated the role of ecological traits, activity patterns, and reproductive pressures in shaping SSD (e.g., *p* < 0.01 for differences in tree vs. ground-dwelling species). This study underscores the complexity of SSD, influenced by ecological, behavioral, and geographic factors. Morphometric indices validated here offer practical tools for future studies, requiring careful application across species, populations, and developmental stages. The results emphasize the ecological and evolutionary importance of body size variation and provide critical data for conservation efforts in Venezuelan anurans.

## INTRODUCTION

In anurans, as in many animal species, differences in body size between males and females are commonly observed, with the magnitude and direction of these differences varying considerably both within and among species (Arak, 1988; López Juri et al., 2018). Sexual size dimorphism (SSD) is hypothesized to be influenced by life history traits such as growth rate (Cohen & Alford, 1993) and age at sexual maturity (Morrison & Hero, 2003). Furthermore, in species with broad geographic distributions, adult body size often exhibits significant phenotypic plasticity (Angilletta et al., 2004; Özdemir et al., 2012). Among anurans, body size is positively correlated with longevity due to their indeterminate growth (Duellman & Trueb, 1994).

Sexual dimorphism in anurans encompasses not only body size but also traits such as coloration, morphology, and behavior. These differences are likely shaped by sexual and/or natural selection (Fairbairn, 1997; Woolbright, 1983). However, the assessment of sexual dimorphism faces two key challenges: (1) measurement error, stemming from factors such as instrument precision, operator variability, or environmental conditions, and (2) the distinction between statistical and biological relevance (Hayek et al., 2001; Hayek & Heyer, 2005).

Three general patterns of sexual size dimorphism are recognized in amphibians: *female-biased* (females larger than males), *male-biased* (males larger than females), and *monomorphic* (both sexes of similar size) (Shine, 1979; Zhang et al., 2014). Shine (1979) reported that approximately 90% of anuran species exhibit female-biased SSD, often attributed to reproductive advantages conferred by larger female body sizes, such as greater egg-carrying capacity (Salthe & Mecham, 1974).

In species with male-biased SSD, males often display agonistic behaviors, enabling them to defend courtship territories or compete for mates, especially when females are larger (Howard, 1981). These males may also exhibit specialized morphological traits, such as lip spines or fang-like projections on the lower jaw (Duellman & Trueb, 1994; Shine, 1979). In addition, body size can play a role in courtship, where larger males may be preferred by females. Therefore, the interaction between competition between males and the choice of females may favor the development of larger body size in males (Tejedo, 1988; Wells, 1979).

The investigation of SSD in anurans provides insights into the biological and ecological factors influencing size variation between sexes. Several hypotheses have been proposed to explain the observed patterns of SSD:

1. **Environmental Influences**: Body size in anurans is shaped by environmental factors such as temperature and resource availability (Narins & Smith, 1986; Teder & Tammaru, 2005). Although Bergmann’s rule suggests a positive relationship between body size and colder temperatures (Liao & Lu, 2010; Olalla-Tárraga & Rodríguez, 2007), some studies have reported weak correlations between environmental variables and body size (Goldberg et al., 2018).
2. **Behavior and Territoriality**: Larger males may achieve greater reproductive success through physical contests or displays, which often occur in territorial or polygynous species (Howard, 1984; Katsikaros & Shine, 1997; Shine, 1979; Székely et al., 2004; Wells, 1977). However, even in smaller species with female-biased SSD, male-male competition can be frequent (Costa et al., 2010; Haddad, 1991).
3. **Reproductive Trade-offs**: Females may prioritize somatic growth, delaying sexual maturity to achieve larger sizes and higher fecundity (Iturra Cid et al., 2010; Monnet & Cherry, 2002; ; Shine, 1979). Conversely, larger males often incur higher energetic costs and faster weight loss during the breeding season (Wells, 1978).
4. **Differential Mortality**: Female-biased SSD may arise from higher mortality rates in males (Shine, 1979; Vargas Salinas, 2006), often linked to conspicuous behaviors such as vocalizations that increase predation risk (Angulo, 2006; Duellman & Trueb, 1994; Wells, 2007). These behaviors not only communicate sexual receptivity and location but may also amplify the visibility of larger males (Howard, 1981).

Given the importance of body size in understanding the behavior, population dynamics, and reproductive success of animals, as well as their interactions with the environment, morphometrics, or the quantitative description of shapes, it represents a crucial tool in biological research (Bernal & Clavijo, 2009; Rohlf, 1990). This discipline provides a framework to explain various ecological and evolutionary phenomena that generate morphological differences between individuals, such as diseases, ontogenetic development, and adaptation to local geographical factors (Zelditch et al., 2004). In addition, patterns of body variation between species may reveal adaptive divergence (Fairbairn, 1997; Morrison & Hero, 2003).

Morphometrics, the quantitative analysis of form and structure, is a powerful tool for understanding the ecological and evolutionary processes underlying sexual dimorphism. This approach has been instrumental in explaining morphological variations influenced by factors such as disease, ontogenetic development, and local adaptations (Bernal & Clavijo, 2009; Zelditch et al., 2004). At lower taxonomic levels, morphometric analyses facilitate species classification and the identification of cryptic species (Bickford et al., 2007; Lee, 1982).

Zoometric indices, on the other hand, allow quantifying the phenotypic variability of anatomical proportions and directly determining the relationship between the variables of interest. They can be easily calculated and require simple measurements in the field, with little or no impact on individuals. However, these indices must be validated for each species, population, and sex to ensure applicability across age groups (Băncilă et al., 2010).

In Venezuela, anurans represent approximately 5% of global amphibian diversity, with 370 species reported across 16 families (Barrio-Amorós et al., 2019). More than half of these species are endemic, reflecting the country’s diverse habitats shaped by its unique geographic and climatic conditions (Molina et al., 2009). Body sizes among Venezuelan anurans range from less than one centimeter to over 20 centimeters, particularly in families such as Bufonidae, Hylidae, Leptodactylidae, and Phyllomedusidae. Despite some studies addressing morphological differences in certain species, such as *Aromobates tokuko* (Rojas-Runjaic et al., 2011), *Hyloscirtus japreria* (Rojas-Runjaic et al., 2018), and *Rhinella marina* (Acevedo et al., 2016), comprehensive assessments of SSD in Venezuelan anurans remain scarce.

This study aims to explore variations in body size, body proportions, and sexual dimorphism in Venezuelan anurans, examining their potential associations with geographic distribution, habitat characteristics, and behavioral patterns.

## MATERIALS AND METHODS

### Geographical area

This study focuses on the bioregion known as the **Cordillera de la Costa** in Venezuela. This mountain range extends west to east across the central-northern region of the country and is divided into two distinct physiographic sections, separated by the **Unare River Valley** (Madi et al., 2011). The central section spans the states of Yaracuy, Carabobo, Aragua, Guárico, and Miranda, including the Capital District, while the eastern section primarily encompasses Anzoátegui, Monagas, and Sucre states (Fig. 1). The Cordillera de la Costa reaches a maximum elevation of slightly over 2,500 meters above sea level (masl). The region’s vegetation ranges from savannas to cloud forests. The northern slope is dominated by deciduous mountain forests and tropophilous scrub, while the southern slope supports montane tropical forests between 400 and 700 masl. Above 900 masl, evergreen cloud forests predominate. During the rainy season (April to November), the region receives monthly rainfall of 100–150 mm, with annual totals averaging approximately 1,400 mm. The mean annual temperature is 27 °C at elevations between 800 and 1,000 masl but drops to 6–12 °C in the highest areas (INAMEH, 2021).

**Figure 1.**
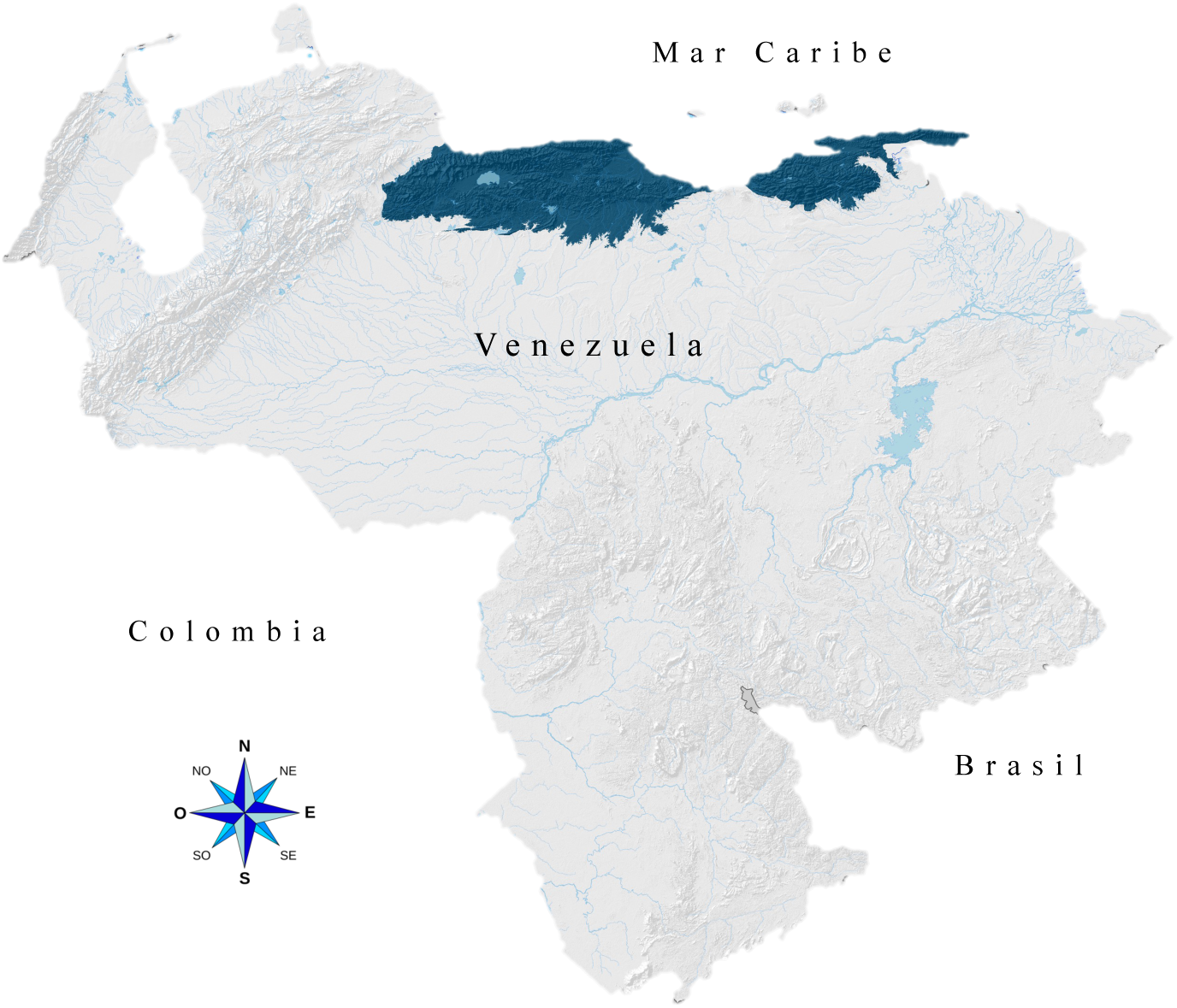
Relative geographic location of the Cordillera de la Costa (shaded dark blue). Source: Madi et al. (2011).

### Taxa considered

The study focuses on anuran species from the central section of the Cordillera de la Costa, representing a subset of the amphibian fauna of this bioregion. This fauna includes 89 species (86 anurans, two caecilians, and one salamander), accounting for 23.2% of Venezuela’s amphibian diversity. Notably, 68.9% of these species are exclusive to the Cordillera de la Costa, and 60% are endemic to Venezuela (Barrio-Amorós et al., 2019). The nomenclature, classification, and systematics of the studied anuran species (Table 1) were verified following Barrio-Amorós et al. (2019).

**Table 1.** List of species studied. ♂♂ = males, ♀♀ = females.

| FAMILY | Sample size (n = 745) |  | Activity<sup>1</sup> | Habitat<sup>1</sup> | Distribution<sup>2</sup> |
| --- | --- | --- | --- | --- | --- |
| Species | ♂♂ | ♀♀ |  |  |  |
| BUFONIDAE |  |  |  |  |  |
| *Atelopus cruciger* (Lichtenstein & Martens, 1856) | 72 | 66 | D | T | Restricted |
| *Rhinella marina* (Linnaeus, 1758) | 25 | 15 | C/N | T | Wide |
| HYLIDAE |  |  |  |  |  |
| *Boana punctata* (Schneider, 1799) | 37 | 76 | N | F | Restricted |
| *Boana xerophylla* (Duméril & Bribon, 1841) | 36 | 15 | N | S | Wide |
| *Dendropsophus luteoocellatus* (Roux, 1927) | 14 | 3 | C/N | S | Restricted |
| *Dendropsophus microcephalus* (Cope, 1886) | 112 | 17 | N | S | Wide |
| *Scarthyla vigilans* (Solano, 1971) | 30 | 26 | D/N** | S** | Wide |
| *Scinax rostratus* (Peters, 1863) | 11 | 8 | N | F, S | Wide |
| LEPTODACTYLIDAE |  |  |  |  |  |
| *Engystomops pustulosus* (Cope, 1864) | 119 | 30 | N | T | Wide |
| *Leptodactylus fuscus* (Schneider, 1799) | 10 | 9 | N | T | Wide |
| PHYLLOMEDUSIDAE |  |  |  |  |  |
| *Phyllomedusa trinitatis* (Meterns, 1926) | 11 | 3 | N | F | Restricted |

<sup>1</sup>Señaris et al. (2014, 2018). <sup>2</sup>Barrio-Amorós et al. (2019). \*\*Rojas-Runjaic et al. (2008).
C = Crepuscular, D = Diurnal, N = Nocturnal / F = Forest, S = Shrubland, T = Terrestrial

Sex determination was conducted initially by examining external secondary sexual characteristics (Kok & Kalamandeen, 2008; Sexton, 1958; Sever & Staub, 2011) and subsequently confirmed through direct observation of gonadal development. Specimens are deposited in the national reference collections of the Rancho Grande Biological Station (EBRG), the La Salle Natural History Museum (MHNLS), and the Central University of Venezuela Museum of Biology (MBUCV), as well as in the teaching collection of the Laboratory of Ecology and Wildlife Management of the Institute of Zoology and Tropical Ecology (LEMFS–IZET) (Appendix I).

### Morphometric data

Morphometric variation was analyzed for 11 anuran species (Table 1).

**A) Linear Morphometry.** Two primary linear measurements were assessed (Watters et al., 2016) (Fig. 2): 1. Snout-vent length (SVL): The linear distance from the tip of the snout to the posterior edge of the cloaca. 2. Abdominal width (AW): The maximum linear distance across the widest point of the body.

**Figure 2.**
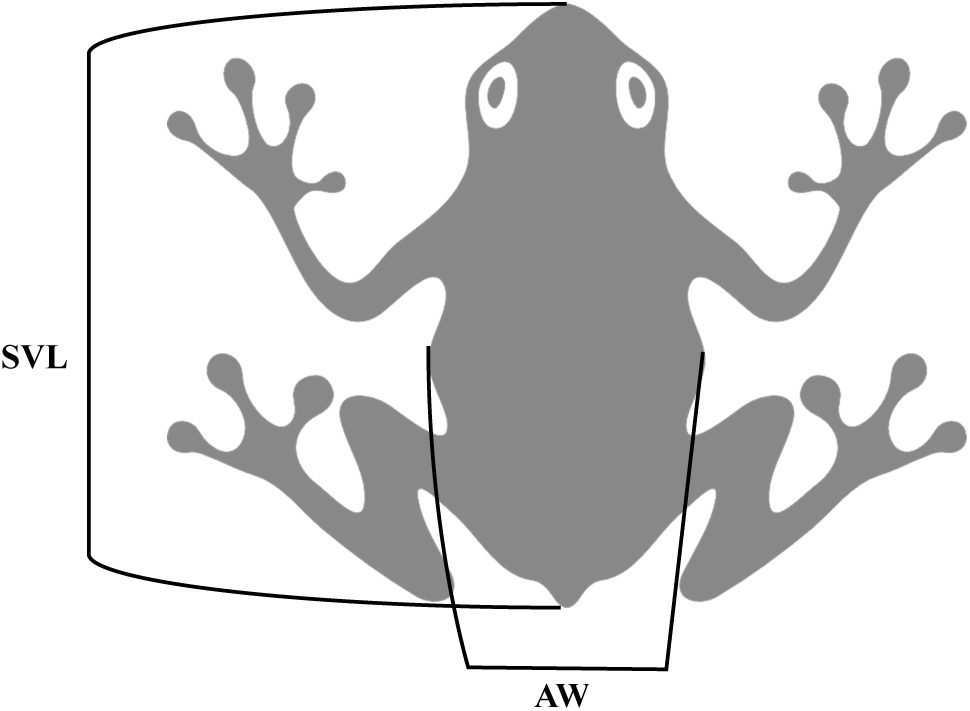
Diagram illustrating measurement points on anurans: SVL (snout-vent length) and AW (abdominal width).

All measurements were recorded using metal Vernier calipers with an accuracy of ±0.5 mm, with each measurement repeated 3–4 times to minimize error. Body sizes were categorized into five classes: *Very small* (< 20 mm), *Small* (20-30 mm), *Medium* (30-60 mm), *Large* (60-200 mm) and, *Very large* (> 200 mm). Classification criteria follow Kok & Kalamandeen (2008).

**B) Zoometric Indices.** Zoometric indices quantify the relationships between morphological variables, facilitating phenotypic comparisons across species and sexes. Two indices were calculated:

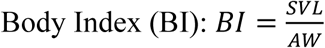

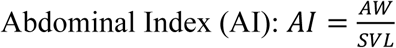

These indices classify individuals into size categories based on length-to-width proportions.

**C) Sexual dimorphism.** Four indices were employed to evaluate sexual dimorphism (SD) in body size:

1. Absolute Difference (*Da*): The absolute difference in mean SVL between sexes:

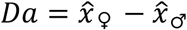 Negative values indicate male-biased dimorphism.
2. Sexual Dimorphism Index (*SDIa*): Following Lovich & Gibbons (1992):

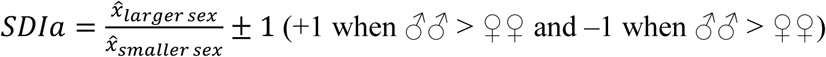 The values are symmetrical around zero (*SDIa* ≈ 0, decreases sexual dimorphism). Values are positive when females are larger and negative when males are larger.
3. Revised Sexual Dimorphism Index (*SDIb*): Modified by Smith (1999):

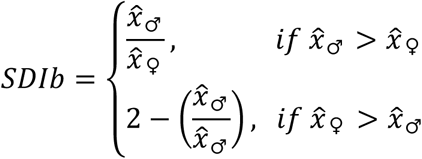 Values are symmetrical around 1(*SDIb* ≈ 1, decreases sexual dimorphism).
4. Percentage Dimorphism Index (*SDIc*): Modified by Angel et al. (2015):

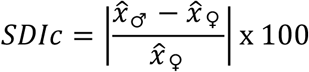 Negative values indicate male-biased dimorphism.

**D) Statistical analysis.** Morphometric data were analyzed separately by sex.

Descriptive statistics were calculated for all variables, with a significance threshold of p ≤ 0.05. Differences in SVL and AW between sexes were tested using the **Mann-Whitney U test**. The variability in coefficients of variation by sex was assessed using the **t-test**. Relationships between zoometric indices and species by sex were quantified using the **correlation coefficient (r)**. Differences in zoometric indices between sexes were evaluated with the **Kruskal-Wallis test**. The influence of habitat type, activity pattern, and geographic distribution on SD was analyzed using **ANOVA**.

All analyses and graphical outputs were performed using PAST 4.0 software (Hammer et al., 2001).

## RESULTS

A total of 745 specimens were analyzed in this study, comprising 64.03% males (n=477) and 35.97% females (n=268).

### Linear morphometry

Table 2 summarizes the descriptive statistics of the snout-vent length (SVL) and abdominal width (AW), stratified by sex for the studied species. Among males, the predominant size category was small (20–30 mm), encompassing *A. cruciger*, *B. punctata, D. microcephalus* and *E. pustulosus*. Medium-sized males (30–60 mm) included *B. xerophylla*, *L. fuscus* and *S. rostratus*, while very small males (<20 mm) were observed in *D. luteoocellatus* and *S. vigilans.* The large size category (60–200 mm) included *P. trinitatis* and *R. marina.* In females, the small category (20–30 mm) comprised *B. punctata, D. luteoocellatus, D. microcephalus, E. pustulosus,* and *S. vigilans.* Large females (60–200 mm) were observed in *B. xerophylla, L. fuscus, P. trinitatis,* and *R. marina,* while medium-sized females (30–60 mm) included *A. cruciger* and *S. rostratus*.

**Table 2.** Descriptive statistics of the snout-vent length^a^ and abdominal width^b^ of the 11 species discriminated by sex. Values in mm. (p ≤ 0.05*).

| Species | Males |  |  |  | Females |  |  |  |  |  |
| --- | --- | --- | --- | --- | --- | --- | --- | --- | --- | --- |
| | Mean ± SD | Min. - Max. (n) | CV | Me | Mean ± SD | Min. - Max. (n) | CV | Me | U | p |
| *A. cruciger* | 28.73 ± 4.31<sup>a</sup> | 17 - 37 (72) | 0.15 | 29 | 38.66 ± 5.36 | 30 - 51 (66) | 0.14 | 37 | 264.0 | 0.000* |
| | 8.99 ± 1.75<sup>b</sup> | 6 - 13 (72) | 0.19 | 9 | 12.26 ± 2.43 | 9 - 18 (66) | 0.2 | 12 | 594.0 | 0.000* |
| *R. marina* | 84.12 ± 22.84<sup>a</sup> | 43 - 120 (25) | 0.27 | 85 | 85.87 ± 28.28 | 42 - 117 (15) | 0.33 | 95 | 169.0 | 0.614 |
| | 46.76 ± 18.44<sup>b</sup> | 17 - 76 (25) | 0.39 | 44 | 48.20 ± 19.65 | 19 - 73 (15) | 0.41 | 51 | 180.5 | 0.856 |
| *B. punctata* | 27.00 ± 7.12<sup>a</sup> | 16 - 36 (37) | 0.26 | 29 | 25.99 ± 7.44 | 16 - 39 (76) | 0.29 | 27 | 1299.0 | 0.514 |
| | 10.19 ± 3.53<sup>b</sup> | 5 - 16 (37) | 0.35 | 10 | 9.11 ± 2.85 | 4 - 15 (76) | 0.31 | 9 | 1161.0 | 0.132 |
| *B. xerophylla* | 55.17 ± 3.62<sup>a</sup> | 47 - 62 (36) | **0.07** | 56 | 60.07 ± 2.87 | 56 - 66 (15) | **0.05** | 60 | 76.0 | 0.000* |
| | 19.31 ± 2.29<sup>b</sup> | 14 - 24 (36) | 0.12 | 20 | 19.6 ± 3.72 | 12 - 25 (15) | 0.19 | 21 | 222.0 | 0.323 |
| *D. luteoocellatus* | 18.50 ± 1.02<sup>a</sup> | 17 - 21 (14) | **0.06** | 19 | 23.00 ± 1.00 | 22 - 24 (3) | **0.04** | 23 | 0.0 | 0.007* |
| | 5.21 ± 0.70<sup>b</sup> | 4 - 7 (14) | 0.13 | 5 | 9.67 ± 0.58 | 9 - 10 (3) | **0.06** | 10 | 0.0 | 0.004* |
| *D. microcephalus* | 20.65 ± 1.70<sup>a</sup> | 17 - 24 (112) | **0.08** | 21 | 23.59 ± 2.83 | 19 - 27 (17) | 0.12 | 24 | 398.5 | 0.000* |
| | 7.20 ± 1.11<sup>b</sup> | 5 - 10 (112) | 0.15 | 7 | 9.18 ± 1.67 | 6 - 11 (17) | 0.18 | 10 | 335.5 | 0.000* |
| *S. vigilans* | 17.53 ± 1.25<sup>a</sup> | 14 - 21 (30) | **0.07** | 18 | 22.27 ± 1.28 | 19 - 24 (26) | **0.06** | 22 | 5.5 | 0.000* |
| | 5.53 ± 0.82<sup>b</sup> | 4 - 7 (30) | 0.15 | 6 | 7.46 ± 1.10 | 6 - 10 (26) | 0.15 | 7 | 60.0 | 0.000* |
| *S. rostratus* | 41.27 ± 2.57<sup>a</sup> | 36 - 44 (11) | **0.06** | 42 | 42.00 ± 2.83 | 37 - 45 (8) | **0.07** | 42 | 36.0 | 0.528 |
| | 16.36 ± 1.86<sup>b</sup> | 13 - 20 (11) | 0.11 | 16 | 16.50 ± 2.98 | 10 - 19 (8) | 0.18 | 43 | 35.5 | 0.502 |
| *E. pustulosus* | 26.85 ± 2.21<sup>a</sup> | 21 - 31 (119) | **0.08** | 27 | 28.57 ± 1.68 | 24 - 32 (30) | **0.06** | 29 | 985.0 | 0.000* |
| | 13.02 ± 2.38<sup>b</sup> | 9 - 17 (119) | 0.18 | 13 | 14.70 ± 1.91 | 10 - 16 (30) | 0.13 | 19 | 1011.5 | 0.000* |
| *L. fuscus* | 54.50 ± 24.76<sup>a</sup> | 28 - 89 (10) | 0.45 | 51 | 61.56 ± 12.32 | 40 - 79 (9) | 0.2 | 63 | 38.0 | 0.595 |
| | 17.80 ± 8.77<sup>b</sup> | 9 - 33 (10) | 0.49 | 15 | 20.67 ± 5.50 | 15 - 31 (9) | 0.27 | 19 | 32.0 | 0.304 |
| *P. trinitatis* | 66.45 ± 3.86<sup>a</sup> | 59 - 74 (11) | **0.06** | 67 | 79.00 ± 1.00 | 78 - 80 (3) | **0.01** | 79 | 0.0 | 0.011* |
| | 18.64 ± 2.20<sup>b</sup> | 15 - 21 (11) | 0.12 | 20 | 23.34 ± 2.89 | 20 - 25 (3) | 0.12 | 25 | 4.0 | 0.056* |

SD = Standard Deviation, Min. = Minimal, Max. = Maximal, (n) = sample size, CV = coefficient of variation (**Bold** = low variation), Me = Median, U = Mann-Whitney, p = significance level.

No significant differences in SVL and AW between males and females were observed for *B. punctata, L. fuscus, R. marina,* and *S. rostratus*. Similarly, there were no significant differences in AW between sexes for *B. xerophylla*. Variation coefficients for both SVL and AW exceeded 10% in most cases, indicating moderate to high variability, but no significant differences were found between sexes (t = 0.315, df = 19, p= 0.378; t = 0.332, df = 19, p = 0.372). Due to insufficient female representation, *D. luteoocellatus* and *P. trinitatis* were excluded from some analyses.

Figure 3 illustrates the distribution of body size values by sex and species using violin plots, highlighting multimodal trends.

**Figure 3.**
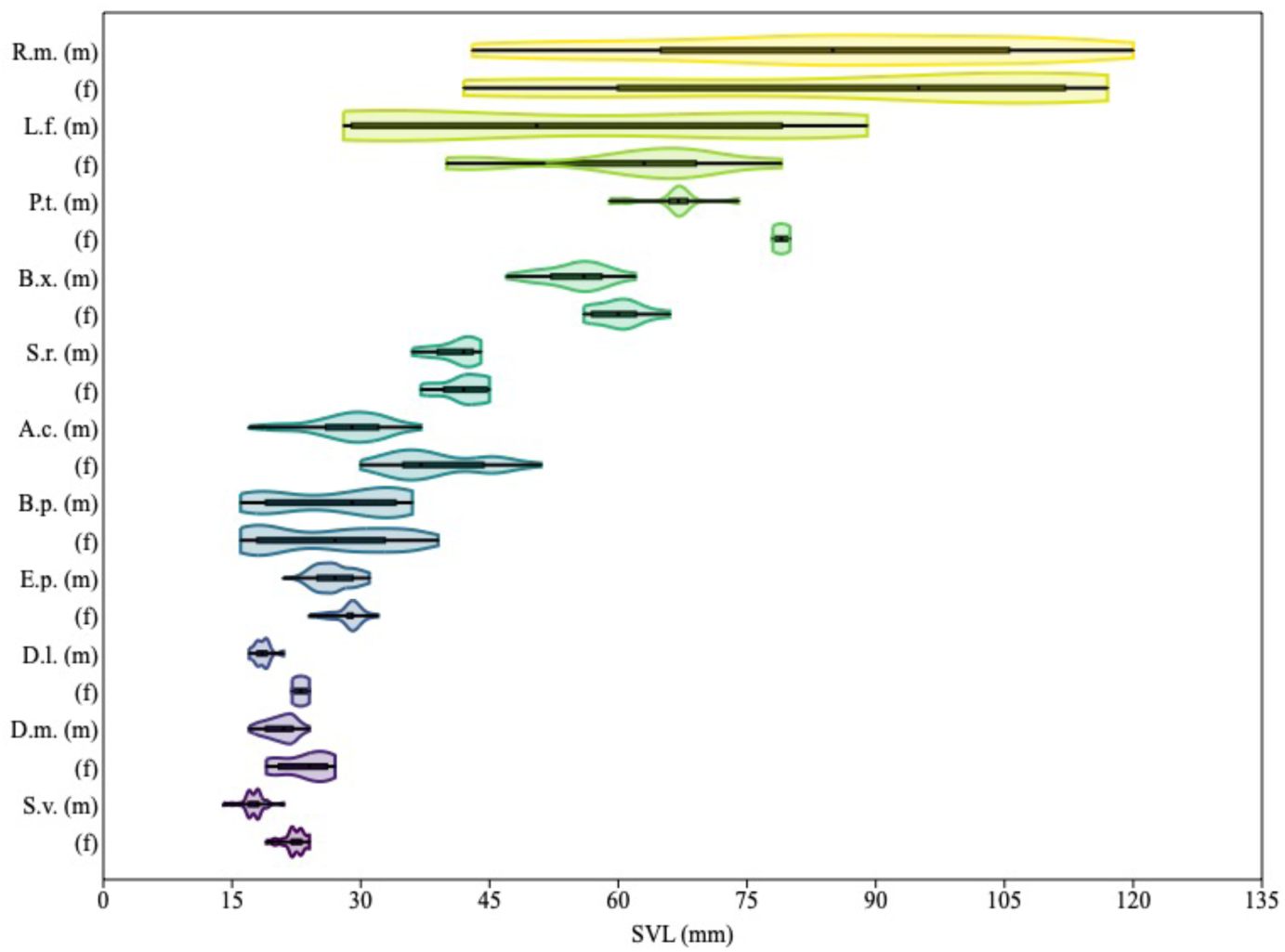
Violin diagram of total body length. The central thick black bar represents the interquartile interval. The thin black lines on the left and right that extend from it represent 95% of the confidence intervals. The colored areas are equivalent to the distribution of the values. A.c. = *A. cruciger*, B.p. = *B. punctata*, B.x. = *B. xerophylla*, D.l. = *D. luteoocellatus*, D.m. = *D. microcephalus*, E.p. = *E. pustulosus*, L.f. = *L. fuscus*, P.t. = *P. trinitatis*, R.m. = *R. marina*, S.v. = *S. vigilans*, S.r. = *S. rostratus*. (m) = males, (f) = females.

Figure 4 presents linear regression analyses of SVL and AW, showing species-specific trends. The determination coefficients (*R^2^*) ranged from negligible (*R^2^* = 0.0004 for *S. rostratus*) to nearly complete (*R^2^* = 0.9094 for *L. fuscus* males).

**Figure 4.**
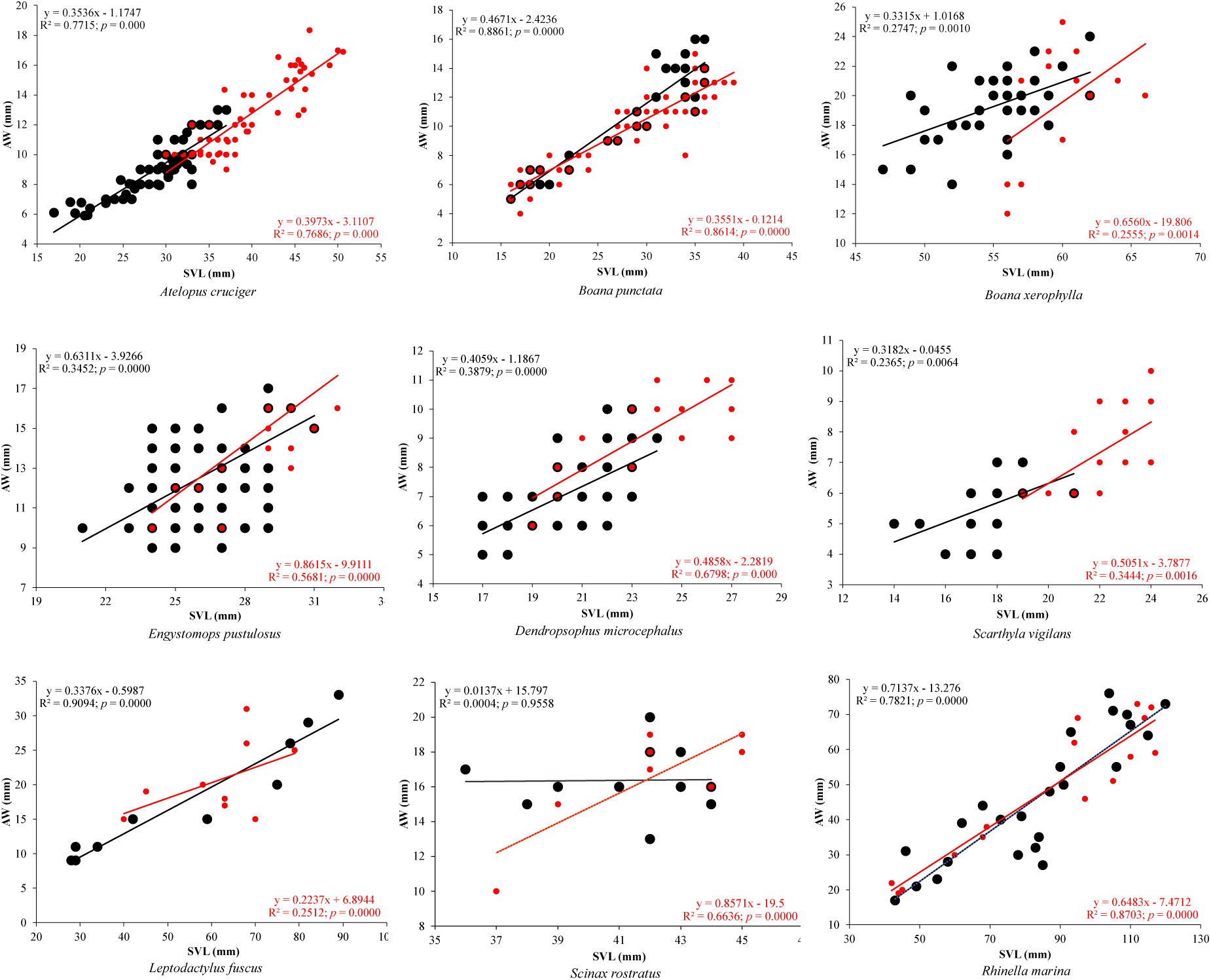
Linear regression of snout-vent length (SVL) and abdominal width (AW) in millimeters (mm). The black color represents the regression of the males. The red color represents the regression of females. The values of the slope, intercept and coefficient of determination are reported.

### Zoometric indices

Tables 3 and 4 detail the body index (BI) and abdominal index (AI) values by sex. High positive correlations were found between the indices across all species. Significant sex-based differences in BI and AI were observed in *D. microcephalus, D. luteoocellatus,* and *E. pustulosus*.

**Table 3.** Body index values (mm) by sex for the 11 species in this study. Kruskal-Wallis (p ≤ 0.05*).

| Species | Males |  |  |  | Females |  |  |  |  |  |
| --- | --- | --- | --- | --- | --- | --- | --- | --- | --- | --- |
| | Mean ± SD | Min. - Max. (n) | Me | r | Mean ± SD | Min. - Max. (n) | Me | r | H | p |
| *A. cruciger* | 3.23 ± 0.29 | 2.64 - 4.13 (72) | 3.23 | 0.997 | 3.20 ± 0.32 | 2.54 - 4.11 (66) | 3.19 | 0.985 | 0.317 | 0.574 |
| *R. marina* | 1.93 ± 0.46 | 1.37 - 3.15 (25) | 1.81 | 0.945 | 1.86 ± 0.28 | 1.38 - 2.32 (15) | 1.91 | 0.987 | 0.020 | 0.888 |
| *B. punctata* | 2.74 ± 0.33 | 2.07 - 3.33 (37) | 2.77 | 0.987 | 2.90 ± 0.38 | 2.14 - 4.25 (76) | 2.83 | 0.948 | 2.563 | 0.109 |
| *B. xerophylla* | 2.89 ± 0.31 | 2.36 - 3.71 (36) | 2.87 | 0.983 | 3.18 ± 0.64 | 2.40 - 4.67 (15) | 3.05 | 0.946 | 1.978 | 0.160 |
| *D. luteocellatus* | 3.59 ± 0.37 | 3.00 - 4.50 (14) | 3.60 | 0.945 | 2.38 ± 0.07 | 2.30 - 2.44 (3) | 2.40 | 0.971 | 7.194 | 0.007* |
| *D. microcephalus* | 2.91 ± 0.35 | 2.20 - 3.67 (112) | 2.86 | 0.983 | 2.61 ± 0.31 | 2.18 - 3.17 (17) | 2.50 | 0.963 | 9.858 | 0.002* |
| *S. rostratus* | 2.55 ± 0.34 | 2.10 - 3.23 (11) | 2.53 | 0.922 | 2.62 ± 0.47 | 2.21 - 3.70 (8) | 2.49 | 0.979 | 0.000 | 1.000 |
| *S. vigilans* | 3.23 ± 0.48 | 2.57 - 4.50 (30) | 3.00 | 0.982 | 3.03 ± 0.35 | 2.40 - 3.67 (26) | 3.14 | 0.853 | 0.932 | 0.334 |
| *E. pustulosus* | 2.12 ± 0.34 | 1.60 - 3.00 (119) | 2.07 | 0.971 | 1.97 ± 0.22 | 1.81 - 2.70 (30) | 1.81 | 0.863 | 4.119 | 0.042* |
| *L. fuscus* | 3.11 ± 0.43 | 2.64 - 3.93 (10) | 3.05 | 0.943 | 3.09 ± 0.77 | 2.19 - 4.67 (9) | 2.90 | 0.962 | 0.327 | 0.568 |
| *P. trinitatis* | 3.60 ± 0.31 | 3.24 - 4.19 (11) | 3.52 | 0.948 | 3.42 ± 0.46 | 3.12 - 3.95 (3) | 3.20 | 0.906 | 1.047 | 0.306 |

SD = Standard Deviation, Min. = Minimal, Max. = Maximal, (n) = Sample size, Me = Median, r = Correlation coefficient, H = Kruskal-Wallis, p = significance level.

**Table 4.** Abdominal index values (mm) by sex for the 11 species in this study. Kruskal-Wallis (p ≤ 0.05*).

| Species | Males |  |  |  | Females |  |  |  |  |  |
| --- | --- | --- | --- | --- | --- | --- | --- | --- | --- | --- |
| | Mean ± SD | Min. - Max. (n) | Me | r | Mean ± SD | Min. - Max. (n) | Me | r | H | p |
| *A. cruciger* | 0.31 ± 0.03 | 0.24 - 0.38 (72) | 0.31 | 0.979 | 0.32 ± 0.03 | 0.24 - 0.39 (66) | 0.31 | 0.990 | 0.233 | 0.630 |
| *R. marina* | 0.55 ± 0.11 | 0.32 - 0.73 (25) | 0.55 | 0.982 | 0.55 ± 0.09 | 0.43 - 0.73 (15) | 0.52 | 0.970 | 0.016 | 0.899 |
| *B. punctata* | 0.37 ± 0.05 | 0.30 - 0.48 (37) | 0.36 | 0.971 | 0.35 ± 0.04 | 0.24 - 0.47 (76) | 0.35 | 0.988 | 2.573 | 0.109 |
| *B. xerophylla* | 0.35 ± 0.04 | 0.27 - 0.42 (36) | 0.35 | 0.992 | 0.33 ± 0.06 | 0.21 - 0.42 (15) | 0.33 | 0.988 | 2.139 | 0.144 |
| *D. luteocellatus* | 0.28 ± 0.03 | 0.22 - 0.33 (14) | 0.28 | 0.960 | 0.42 ± 0.01 | 0.41 - 0.43 (3) | 0.42 | 1.000 | 7.194 | 0.007* |
| *D. microcephalus* | 0.35 ± 0.04 | 0.27 - 0.45 (112) | 0.35 | 0.989 | 0.39 ± 0.04 | 0.32 - 0.46 (17) | 0.40 | 0.972 | 10.580 | 0.001* |
| *S. rostratus* | 0.40 ± 0.05 | 0.31 - 0.48 (11) | 0.39 | 0.963 | 0.39 ± 0.06 | 0.27 - 0.45 (8) | 0.40 | 0.964 | 0.000 | 1.000 |
| *S. vigilans* | 0.32 ± 0.04 | 0.22 - 0.39 (30) | 0.34 | 0.993 | 0.34 ± 0.04 | 0.27 - 0.42 (26) | 0.32 | 0.927 | 1.290 | 0.256 |
| *E. pustulosus* | 0.48 ± 0.07 | 0.33 - 0.63 (119) | 0.48 | 0.982 | 0.51 ± 0.07 | 0.37 - 0.55 (30) | 0.55 | 0.882 | 4.247 | 0.039* |
| *L. fuscus* | 0.33 ± 0.04 | 0.25 - 0.38 (10) | 0.33 | 0.976 | 0.34 ± 0.08 | 0.21 - 0.46 (9) | 0.34 | 0.995 | 0.378 | 0.539 |
| *P. trinitatis* | 0.28 ± 0.02 | 0.24 - 0.31 (11) | 0.28 | 0.954 | 0.29 ± 0.04 | 0.25 - 0.32 (3) | 0.31 | 0.925 | 1.057 | 0.304 |

SD = Standard Deviation, Min. = Minimal, Max. = Maximal, (n) = Sample size, Me = Median, r = Correlation coefficient, H = Kruskal-Wallis, p = significance level.

Based on the data on minimum (Min.) and maximum (Max.) values presented in Tables 3 and 4 and considering the largest number of individuals in each interval, three height classes were established for both the BI and the AI. BI values (1.37–4.67 mm) categorized individuals into three classes: brevilinear (1.37–2.47), mesolinear (2.47–3.57), and longilinear (3.57–4.67). Most species were concentrated in Classes I and II, representing 92.35% (n = 688) of the sample. Class I excludes *A. cruciger* and *P. trinitatis*. In class III, the absent species are *E. pustulosa* and *R. marina.* In contrast, AI values (0.21–0.73 mm) defined three classes: narrow (0.21–0.38), medium (0.38–0.56), and wide (0.56–0.73). Classes I and II accounted for 94.36% (n=703) of individuals. No individuals of *P. trinitatis* in class II were found. Only *E. pustulosa* and *R. marina* were present in class III (n < 25). Figure 5 presents box-and-whisker plots, visualizing sex-specific variations in both indices.

**Figure 5.**
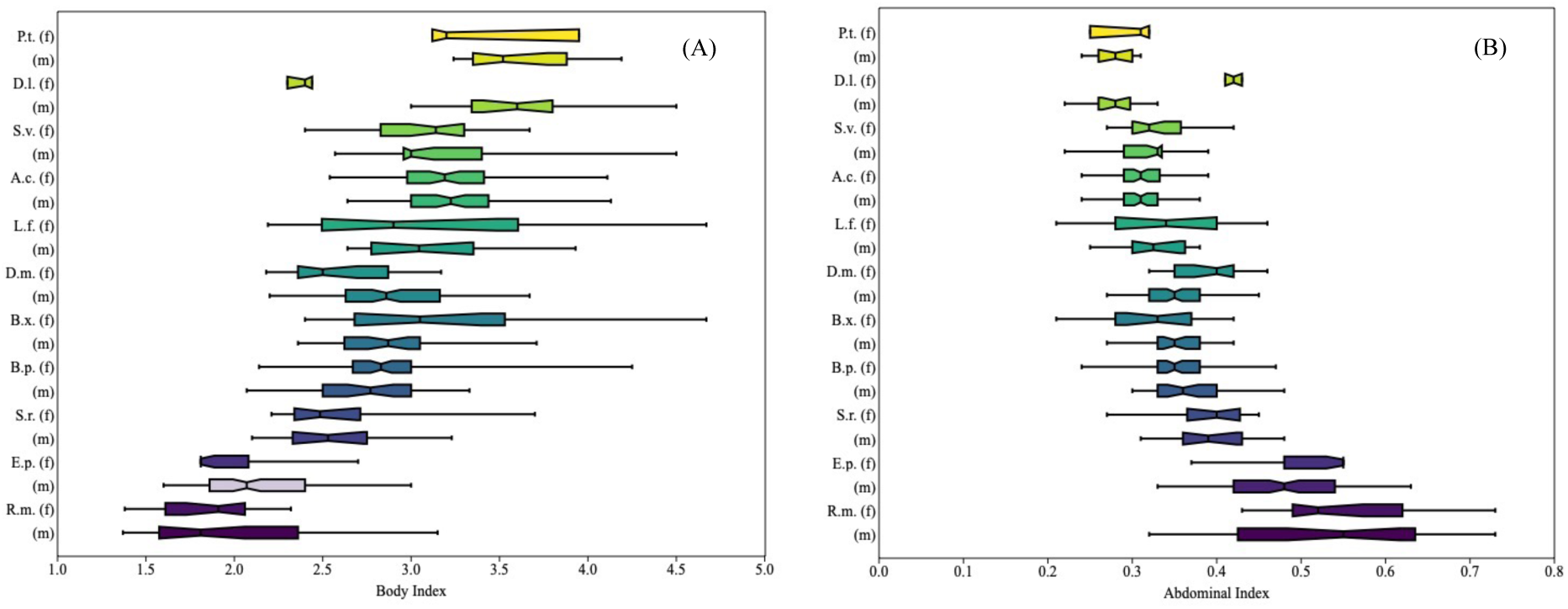
(A) Body index, (B) Abdominal index. The vertical black line inside the boxes corresponds to the median. The colored areas are equivalent to the upper and lower quartiles. The left and right horizontal lines range standard deviation values. (h) = females, (m) = males.

### Sexual dimorphism

All indices of sexual dimorphism (*Da*, *SDIa*, *SDIb*, and *SDIc*) exhibited high positive correlations (r = 0.8289–0.9948). Table 5 summarizes SDI values for each species obtained in this study, while Figure 6 illustrates the direction and magnitude of SDI.

**Figure 6.**
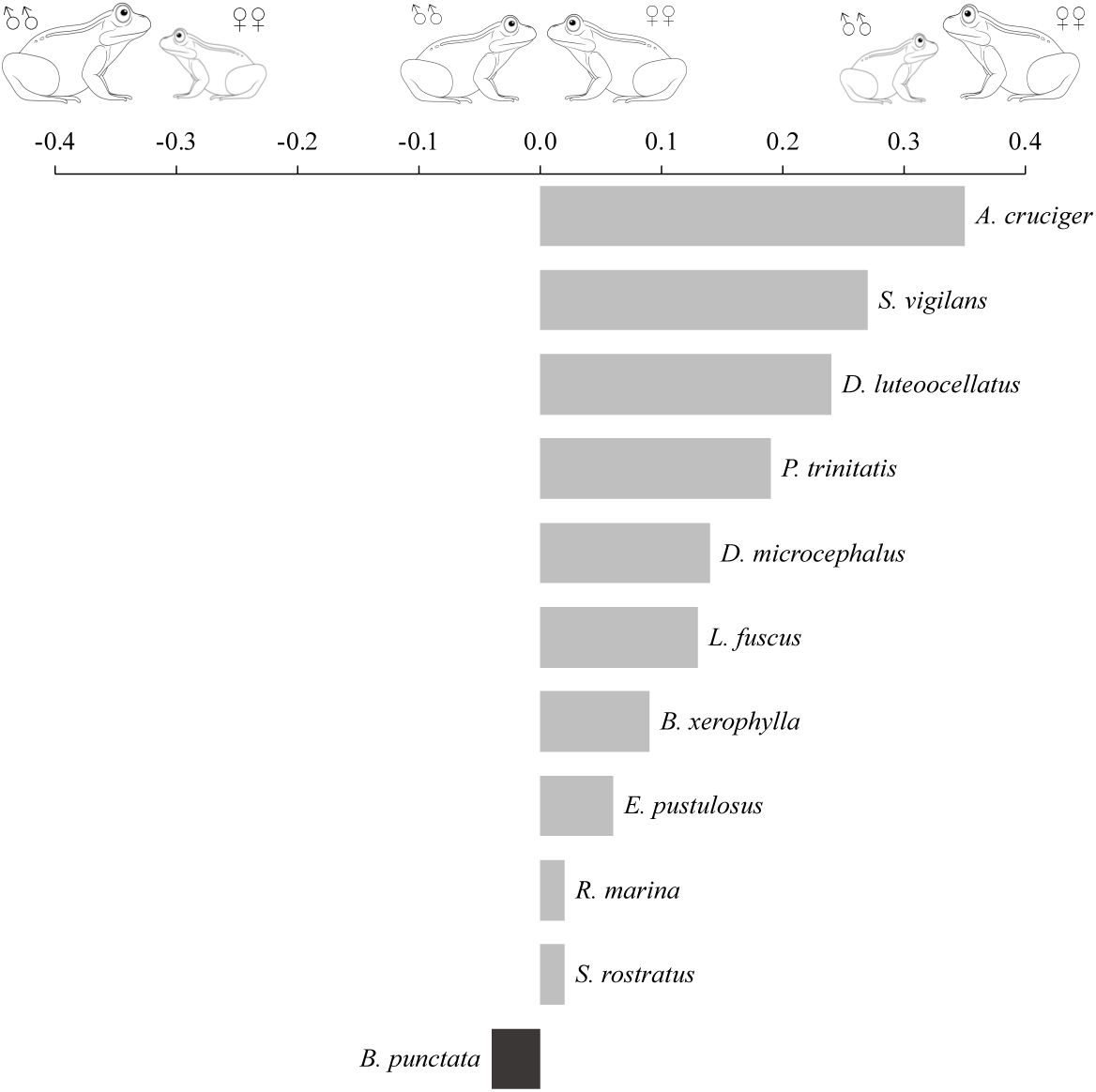
Magnitude and direction of the sexual dimorphism index. The grayscale bars represent the variation of sexual dimorphism based on the dataset. Negative values indicate male bias, while positive values represent female bias. ♂♂ = males, ♀♀ = females.

**Table 5.**
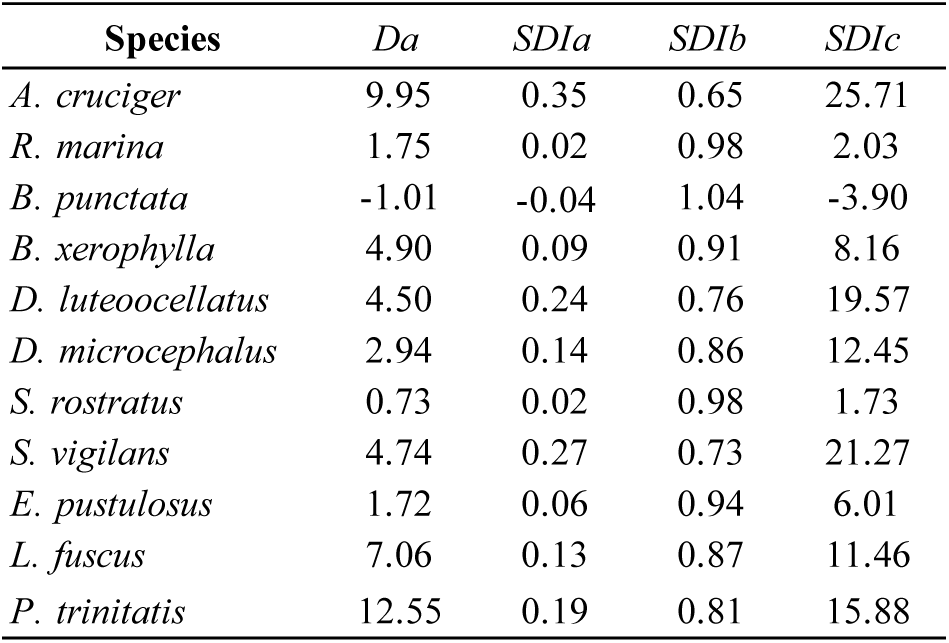
Sex Dimorphism Index (SDI) Values.

| Species | $Da$ | $SDIa$ | $SDIb$ | $SDIc$ |
| --- | --- | --- | --- | --- |
| *A. cruciger* | 9.95 | 0.35 | 0.65 | 25.71 |
| *R. marina* | 1.75 | 0.02 | 0.98 | 2.03 |
| *B. punctata* | -1.01 | -0.04 | 1.04 | -3.90 |
| *B. xerophylla* | 4.90 | 0.09 | 0.91 | 8.16 |
| *D. luteoocellatus* | 4.50 | 0.24 | 0.76 | 19.57 |
| *D. microcephalus* | 2.94 | 0.14 | 0.86 | 12.45 |
| *S. rostratus* | 0.73 | 0.02 | 0.98 | 1.73 |
| *S. vigilans* | 4.74 | 0.27 | 0.73 | 21.27 |
| *E. pustulosus* | 1.72 | 0.06 | 0.94 | 6.01 |
| *L. fuscus* | 7.06 | 0.13 | 0.87 | 11.46 |
| *P. trinitatis* | 12.55 | 0.19 | 0.81 | 15.88 |

Species such as *B. punctata* exhibited male-biased dimorphism, whereas *A. cruciger, B. xerophylla, D. luteoocellatus, D. microcephalus, E. pustulosus, L. fuscus, P. trinitatis, R. marina, S. rostratus,* and *S. vigilans* demonstrated female-biased dimorphism. Notably, *A. cruciger* exhibiting the largest deviation from zero, indicating pronounced female bias. In contrast, *B. punctata* showed a male-biased pattern.

Variance analyses revealed significant differences in sexual dimorphism associated with activity patterns (F_3,40_ = 6.301, p = 0.001), but not with habitat type (F_3,40_ = 0.842, p = 0.479) or geographic distribution (F_1,20_ = 2.134, p = 0.160) Figure 7 highlights these relationships.

**Figure 7.**
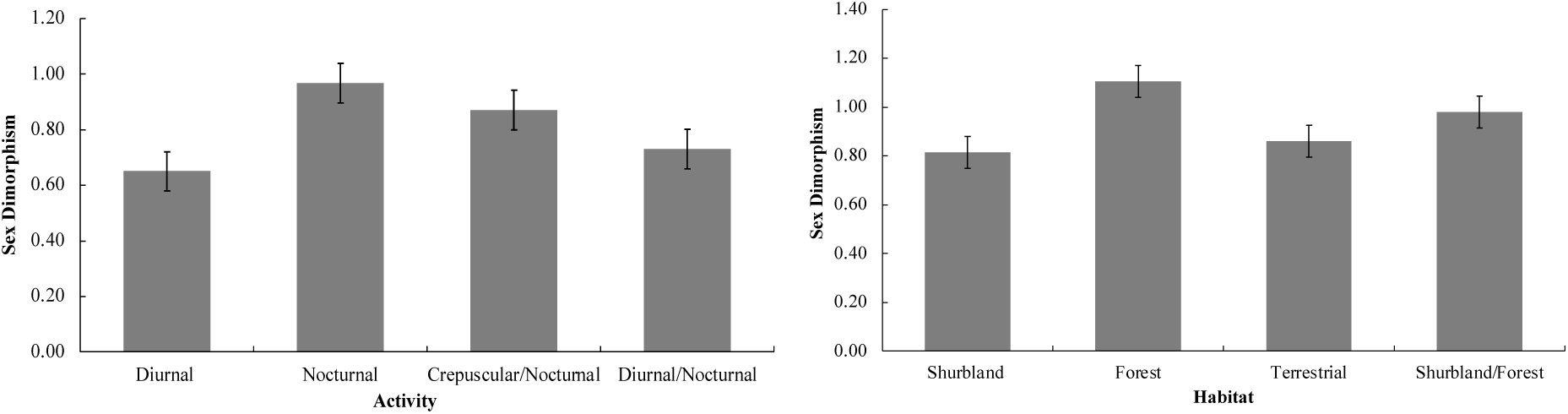
Sexual dimorphism across habitat types and activity patterns for 11 species.

## DISCUSSION

This study provides robust evidence of significant intra- and interspecific differences in snout–vent length (SVL) and abdominal width (AW) among anuran species from Venezuela. Seven of the 11 species analyzed exhibited clear sexual dimorphism, with females typically larger than males. These findings not only corroborate but also refine previously established patterns of sexual size dimorphism (SSD) in anurans. Moreover, this is the first study to report AW values as reference points, underscoring the novelty and relevance of these morphometric data for ecological and evolutionary research.

Notably, four species—*Boana punctata*, *Leptodactylus fuscus*, *Rhinella marina*, and *Scinax rostratus*—did not exhibit significant differences in SVL or AW between sexes. The lack of SSD in these species challenges the expectation of widespread female-biased dimorphism in anurans and raises questions about the ecological and evolutionary mechanisms shaping body size. In terms of average and absolute values, females were found to be larger in all species, confirming a pattern widely observed in previous research (Baraquet et al., 2018; Bosch & Márquez, 1996; Chiapero et al., 2019; Feng et al., 2015; Liao & Lu, 2010; Özdemir et al., 2012; Pesarakloo et al., 2018). While female-biased SSD predominates across amphibians, exceptions are not rare, as noted by Shine (1979) and Han and Fu (2013), who reported male-biased SSD in only 3–4% of species studied. These deviations suggest that SSD is highly context-dependent, shaped by species-specific ecological pressures, reproductive strategies, and life history traits.

Across species, females were consistently larger than males, with rare exceptions such as *B. punctata*. This aligns with findings in other taxa (Baraquet et al., 2018; Bosch & Márquez, 1996; Chiapero et al., 2019; Feng et al., 2015; Liao & Lu, 2010), emphasizing a well-documented link between female size and fecundity. Larger females can allocate more energy to reproduction, producing more or larger eggs—a selective advantage in environments with variable resource availability. However, the absence of significant SSD in some species, including *R. marina* and *L. fuscus*, points to the potential role of additional factors such as sample composition, environmental variability, or stage-specific mortality rates.

The findings of this study confirm and expand upon previously reported differences in body size among anuran species (Castro, 2015; Fonseca et al., 2017; Señaris et al., 2014, 2018; Solano, 1987; Viña, 2015). However, the observed mean snout–vent length (SVL) values for both sexes were lower than those reported for *A. cruciger* by Castro (2015) and Señaris et al. (2018); for *B. punctata* by Señaris et al. (2014); for *D. microcephalus* by Fonseca et al. (2017) and Señaris et al. (2018); and for *D. luteoocellatus* and *S. rostratus* by Señaris et al. (2018). Conversely, the SVL values recorded for *S. vigilans* were slightly higher than those noted by Fonseca et al. (2017).

Interestingly, the SVL of both sexes in *Boana xerophylla* fell within the range reported by Señaris et al. (2018), contrasting with the pronounced morphometric variation observed in the congeneric species *Boana cordobae* (Barrio, 1965) across different geographical regions in Argentina (Baraquet et al., 2012, 2018).

Geographic variation further complicates interpretations of body size and SSD. For instance, *E. pustulosus* exhibited SVL values slightly above those reported for northern Venezuela (Viña, 2015) but lower than those from the Caracas Valley (Señaris et al., 2018). This pattern of variation between geographic locations has also been noted for the species by Atencia (2017) and Atencia et al. (2020) in three locations in northern Colombia.

Likewise, regarding the absence of variation in relation to the size found in other species of anurans present in Venezuela, Molina (2004) points out in *Pleuroderma brachyops* that males were larger than females, but the differences were not statistically significant (t= 2.05; p= 0.052; n= 12).

Similarly, *L. fuscus* showed higher SVL values than reported by Lucas et al. (2008), Maragno & Cechin (2009), Martins (1988), Solano (1987), and Señaris et al. (2018). These discrepancies underscore the influence of local environmental conditions, such as temperature gradients, precipitation patterns, and resource availability, on morphometric traits. Previous studies (e.g., Goldberg et al., 2018) have highlighted the role of climatic and geographic variables in driving body size variation and SSD, but the weak associations reported in some cases suggest the interplay of additional, unmeasured factors.

Within the genus *Leptodactylus*, Hayek and Heyer (2005) highlighted a discrepancy between statistical and biological significance for various morphological traits across species. These authors reported that in a broad sample (n = 442) of *L. fuscus* spanning populations from Panama to Argentina, male snout-vent length (SVL) ranged from 32.4 mm to 55.3 mm, while female SVL ranged from 32.2 mm to 56.3 mm, with no statistically significant differences between sexes. However, a smaller sample (n = 206) from Porto Velho, Brazil, revealed narrower SVL ranges for males (34.2–43.7 mm) and females (34.3–44.2 mm), with significant differences (p = 0.009). In the present study, *L. fuscus* exhibited an average male SVL of 54.5 mm and an average female SVL of 61.56 mm, but these differences were not statistically significant (p = 0.595). These findings suggest that geographic variation in SVL exceeds the range of intrapopulation variation, emphasizing the importance of localized population analyses in understanding morphometric patterns.

In *Rhinella marina*, females typically exhibit a significantly greater snout-vent length (SVL) than males, as reported by several studies (Lee, 1982, 2001; Monnet and Cherry, 2002; Señaris et al., 2014, 2018). However, the results of this study reveal no significant differences in SVL between sexes, indicating a monomorphic or unbiased pattern of sexual dimorphism. Similarly, Arantes et al. (2015) reported significant differences in body size for the congeneric species *Rhinella rubescens* and *R. schneideri* in Brazil’s Cerrado, where males were larger than females. These discrepancies may stem from differences in sample size or the inclusion of younger individuals that have not yet reached full adult size in the analysis.

Variation in body size is a common phenomenon across many animal species (Schäuble, 2004; Silva et al., 2008). However, several studies have highlighted significant differences in morphometric measurements depending on whether the specimens are measured as living or preserved organisms (Bernal & Clavijo, 2009; Hayek & Heyer, 2005; Schäuble, 2004). Additionally, discrepancies in measurements are often amplified when multiple observers are involved, potentially introducing variability. In this study, all measurements were conducted by a single observer under standardized conditions, minimizing potential biases and ensuring consistency in the dataset.

In anurans, body condition indices are widely used to assess the relationship between body mass and a linear body measurement (Bancila et al., 2010; Vera-Candioti et al., 2019), as well as the degree of distribution or symmetry of body fat coverage by visual estimation (Jayson et al., 2018). These indices are considered indicators of environmental stress, health and physical fitness of the individual, and usually offer consistent results and reflect representative biological trends.

Morphometric indices, such as those used here, offer valuable tools for assessing intra- and interspecific variation in body size. Their ease of application and minimal impact on individuals make them particularly suited for field studies. However, unvalidated indices can lead to misleading conclusions. Researchers are urged to rigorously test the applicability of these indices across populations, age groups, and ecological contexts to ensure robust interpretations.

The results in this study are compatible with what was pointed out by Santini et al. (2018) and show that directional asymmetry is not particularly rare, and its occurrence can vary within a taxonomic family or between the habitat preferences of species. These differences suggest that tree species are longer and narrower animals than terrestrial species. In tree frogs, longer body length is often associated with longer legs, and better locomotor performance (Bijma et al., 2016). Therefore, body size probably improves the possibility of movement in the tree canopy. These features are probably of minor importance to terrestrial species.

Activity patterns and habitat preferences emerged as significant predictors of SSD. Terrestrial species, often characterized by male territoriality, displayed a tendency toward male-biased SSD, consistent with hypotheses linking larger male size to competitive advantages in mate acquisition. In contrast, arboreal species showed stronger female-biased SSD, likely reflecting the locomotor and reproductive demands of these environments. These findings align with Bijma et al. (2016), who demonstrated that arboreal anurans benefit from elongated body proportions and enhanced locomotor performance, traits often more pronounced in females.

The body size of the species analyzed in this study showed a clear relationship with their patterns of sexual dimorphism: smaller-bodied species exhibited a female-biased dimorphism, whereas larger-bodied species displayed a male-biased pattern. This trend aligns with *Rensch’s Rule*, which posits that sexual dimorphism increases with body size when males are larger, but decreases when females are larger (Fairbairn, 1997). Among the species studied, the most pronounced size variations were observed in females of *A. cruciger*, *B. xerophylla*, *D. luteoocellatus*, *L. fuscus*, *P. trinitatis*, and *S. vigilans*, with differences ranging between 4 and 15 mm. In contrast, smaller absolute size variations (1–3 mm) were detected in *D. microcephalus*, *E. pustulosus*, *R. marina*, and *S. rostratus*. Notably, males and females of *S. rostratus* were nearly identical in size, reflecting minimal sexual dimorphism. Conversely, *B. punctata* males, despite being larger, exhibited only slight size differences compared to females, with a variation of approximately 1 mm.

The sexual dimorphism indices in this study reveal distinct patterns across species. For *B. xerophylla*, *E. pustulosus*, *R. marina*, and *S. rostratus*, the values of *SDIa* (> 0.00 < 0.10) and *SDIb* (> 0.90 < 1.00) suggest a male-biased trend. In contrast, species such as *D. microcephalus*, *L. fuscus*, and *P. trinitatis* exhibit a slight female-biased pattern, reflected in *SDIa* values (> 0.10 < 0.20) and *SDIb* values (> 0.80 < 0.90). A more pronounced female bias is observed in *A. cruciger*, *D. luteoocellatus*, and *S. vigilans*, characterized by *SDIa* values (> 0.20 < 0.40) and *SIDb* values (> 0.60 < 0.80).

Additionally, the lack of significant differences in SVL and AW between males and females of *B. punctata*, *L. fuscus*, *R. marina*, and *S. rostratus* reflects the absence of marked sexual dimorphism in these species. For *L. fuscus*, this absence of dimorphism has been previously documented in populations from Brazil and Venezuela (Hayek & Heyer, 2005; Maragno & Cechin, 2009; Solano, 1987).

As highlighted in the introduction and supported by the results of this study, several factors may explain the varying levels of sexual dimorphism observed in the analyzed anuran species. While the effect of temperature gradients was not directly evaluated here, the diversity of habitat types and the broad geographical distribution of the species studied—both longitudinal and altitudinal—do not appear to influence the observed variation in body size.

Morrison and Hero (2003) suggested that species inhabiting cooler environments grow more slowly, taking longer to reach asymptotic body size and sexual maturity compared to those exposed to higher temperatures. However, a contrasting perspective is offered by Goldberg et al. (2018), who studied *Scinax fuscovarius* across three geographically distinct regions in Argentina, Brazil, and Paraguay. Their findings indicate a weak correlation between environmental variables and both body size variation and sexual dimorphism. They observed that body size showed greater variation with longitude than latitude and was positively correlated with rainfall seasonality, while populations in regions with lower annual rainfall exhibited greater degrees of sexual dimorphism. Notably, mean annual temperature and productivity did not significantly influence body size or dimorphism in either sex. Additional, suggest that yet unidentified factors likely drive the geographical variation observed in *S. fuscovarius*.

In this study, sexual dimorphism was found to vary significantly in relation to activity patterns. Diurnal and nocturnal terrestrial species exhibit territorial defense behaviors, and in these species, males are as large as or larger than females. In contrast, arboreal species display a pattern where males are smaller than females. Consequently, terrestrial species are expected to exhibit greater variation in sexual dimorphism compared to arboreal species.

Heyer (1978) proposed a hypothesis regarding *Leptodactylus fuscus*, linking the construction of underground bridal chambers or burrows to male-biased sexual dimorphism. Males of species within this group are known to construct burrows using their snouts (Prado et al., 2002; Reading & Jofre, 2003). This behavior has been associated with the observation that males possess longer heads than females, likely as an adaptation to this specialized reproductive activity (Heyer, 1978; Ponssa & Barrionuevo, 2012). These findings suggest that sexual dimorphism in certain species may be driven not only by ecological and behavioral factors such as territoriality but also by specific reproductive strategies and habitat-related adaptations.

The theory posits that females allocate more energy toward somatic growth and delay sexual maturity to achieve larger body sizes, which are associated with increased fecundity (Iturra Cid et al., 2010; Monnet & Cherry, 2002; Shine, 1979). Conversely, during the breeding season, larger males in some species experience faster body weight loss compared to their smaller counterparts (Wells, 1978), potentially influencing patterns of sexual dimorphism.

Female-biased dimorphism is likely influenced by higher mortality rates in males (Shine, 1979; Vargas Salinas, 2006). Reduced male survival allows females a longer period to grow, leading to larger body sizes. Male anurans display one of the most conspicuous reproductive behaviors—vocalization or calling (Angulo, 2006; Duellman & Trueb, 1994). This behavior not only signals sexual receptivity, spatial location, and individual size but also increases predation risk (Howard, 1981; Wells, 2007). Larger males, by producing more prominent calls, may be more detectable to predators compared to smaller males. This predation pressure, combined with energetic costs, may contribute to the observed dimorphism in body size between sexes in certain species.

In conclusion, this study reveals distinct patterns of sexual dimorphism and interspecific variation in the body size of anuran species in Venezuela. Our findings highlight the importance of considering these morphological traits in ecological and evolutionary research on anurans, as well as in biodiversity conservation efforts across the region. It is essential to note that the data analyzed pertain to sexually mature individuals, with maturity confirmed by evaluating the degree of gonadal development.

The results confirm several key points: (1) general patterns of body size variation show greater size in females compared to males; (2) the relationship between SVL and AW varies significantly across species, influenced by factors such as activity patterns, habitat types, and geographic distribution; (3) body size differences reflect varying degrees of sexual dimorphism; and (4) while the absence of sexual dimorphism in *L. fuscus*, *R. marina*, and *S. rostratus* is clear, further investigation is needed for *D. luteoocellatus* and *P. trinitatis*, as the limited representation of females in this study warrants additional data for more robust conclusions.

## ACKNOWLEDGEMENTS

To F. Machado Stredel for the suggestions and options in the statistical analysis. To the anonymous reviewers who with their comments helped to improve the quality of the manuscript. To Cesar Molina Rodríguez (1960 – 2015) This work is a pending debt to a herpetologist, ecologist, university professor and friend.

## ETHICAL STATEMENT

To carry out this study, access to the biological material of each institution was duly authorized by the administrative managers of each center. All handling protocols followed the principles of good practices required in the care, management and conservation of biological collections. Every effort was made to use only the minimum number of animals necessary to produce reliable scientific data.

## APPENDIX I

*Atelopus cruciger*. **EBRG:** 10; 11; 12; 89; 90; 91; 92; 93; 94; 95; 96; 1062; 1208; 1209; 1210; 1211; 1212; 1213; 1214; 1606; 3968; 3969; 4020; 4021; 4033; 4034; 5097; 5510; 5511. **MBUCV**: 156a; 156b; 156c; 156d; 159; 620a; 620b; 721a; 721b; 722a; 722b; 723a; 723b; 723c; 723d; 724a; 724b; 725a; 725a; 725b; 725b; 725c; 725c; 725d; 725d; 725e; 725f; 725g; 725h; 725i; 726a; 726b; 726c; 726d; 726e; 726f; 726g; 726h; 726i; 726j; 726k; s/nc1; s/nc2; s/nc3. **MHNLS:** 176; 177; 178; 182; 793; 1076; 1077; 1179; 1321; 1322; 1704; 1705; 1706; 1907; 1909; 1910; 1922; 1968; 1969; 2432; 2691; 4327; 4328; 4424; 5542; 5543; 5545; 5547; 5744; 6591; 6594; 6596; 6597; 6597; 6597; 598; 6599; 6600; 6603; 6604; 6606; 6828; 6829; 6864; 6865; 7522; 7525; 7526; 7528; 7529; 7540; 7541; 7542; 7543; 7549; 7551; 7553; 7555; 8453; 8454; 8456; 8458; 11944; 11945; 1306a; 1416a; 1416b.

*Boana punctata*. **LEMFS – IZET**: 145; 166; 167; 168; 169; 170; 171; 172; 173; 253; 254; 255; 256; 268; 271; 272; 273; 274; 275; 276; 277; 278; 279; 280; 281; 282; 283; 284; 285; 286; 286a; 287; 288; 289; 290; 290; 290; 1; 292; 293; 294; 295; 296; 297; 298; 299; 300; 301; 302; 303; 304; 305; 306; 307; 364; 365; 366; 367; 368; 369; 597a; 598; 599; 600; 601; 602; 603; 604; 605; 606; 607; 608; 609; 610; 611; 612; 613; 614; 615; 616; 617; 618; 619; 620; 621; 622; 623; 624; 625; 626; 627; 628; 629; 630; 631; 632; 633; 634; 635; 636; 637; 638; 639; 640; 641; 642; 643; 644; 645; 646; 647; 648; 649; 650; 651.

*Boana xerophylla*. **LEMFS – IZET**: 111; 144; 148; 161; 162; 203; 204; 205; 212; 213; 214; 215; 216; 217; 218; 219; 257; 258; 259; 260; 261; 262; 263; 264; 265; 266; 267; 621; 622; 623; 624; 625; 626; 627; 628; 629; 630; 631; 632; 633; 634; 635; 636; 637; 638; 639; 640; 641; 642; 643; 644.

*Dendropsophus luteoocellatus*. **LEMFS – IZET**: 604; 605; 606; 607; 608; 609; 610; 611; 612; 613; 614; 615; 616; 617; 618; 619; 620.

*Dendropsophus microcephalus*. **LEMFS – IZET**: 370; 371; 372; 373; 374; 375; 376; 377; 378; 379; 380; 381; 382; 383; 384; 385; 386; 387; 388; 389; 390; 391; 392; 393; 394; 395; 396; 397; 398; 399; 400; 401; 402; 403; 404; 404; 5; 406; 407; 408; 409; 410; 411; 412; 413; 414; 415; 416; 417; 418; 419; 420; 421; 422; 423; 424; 425; 426; 427; 428; 429; 430; 431; 432; 433; 434; 435; 436; 437; 438; 439; 440; 441; 442; 443; 444; 445; 446; 447; 448; 449; 450; 451; 452; 453; 454; 455; 456; 457; 458; 459; 460; 533.

*Engystomops pustulosus*. **LEMFS – IZET**: 193; 194; 195; 196; 197; 198; 199; 200; 220; 221; 222; 223; 224; 225; 226; 227; 228; 229; 230; 329; 330; 331; 332; 333; 334; 335; 336; 337; 338; 339; 340; 341; 342; 343; 344; 344; 344; 344 5; 346; 347; 348; 349; 350; 351; 464; 465; 466; 467; 468; 469; 470; 471; 472; 473; 474; 475; 476; 477; 478; 479; 480; 481; 482; 483; 484; 485; 486; 487; 488; 489; 490; 491; 492; 493; 494; 495; 496; 497; 498; 499; 500; 501; 502; 503; 504; 505; 506; 507; 508; 509; 510; 512; 513; 514; 515; 516; 517; 518; 519; 520; 521; 522; 523; 524; 525; 526; 527; 528; 529; 530; 531; 532; 533; 534; 535; 536; 537; 538; 539; 540; 541; 542; 543; 544; 545; 546; 547; 548; 549; 550; 551; 552; 553; 554; 556; 557; 558; 559; 560; 561; 562; 563; 564; 565; 566; 567; 568; 569; 570.

*Leptodactylus fuscus*. **LEMFS – IZET**: 143; 146; 157; 159; 160; 233; 234; 235; 236; 237; 238; 239; 240; 241; 532; 533; 569; 595; 596.

*Phyllomedusa trinitatis*. **LEMFS – IZET**: 149; 201; 202; 242; 243; 244; 245a; 245b; 246; 248; 570; 571; 572; 573.

*Rhinella marina*. **LEMFS – IZET**: 002; 013; 056; 141; 142; 147; 150; 151; 152; 153; 154; 155; 156; 206; 207; 208; 209; 210; 211; 231; 232; 521; 522; 523; 524; 525; 526; 527; 528; 529; 530; 531; 575; 576; 577; 577; 578a; 578b; 579a; 579b; s/n.

*Scinax rostratus*. **LEMFS – IZET**: 153; 162; 163; 164; 165; 247; 249; 250; 251; 252; 580; 581; 582; 583; 584; 585a; 585b; 586; 587.

*Scarthyla vigilans*. **LEMFS – IZET**: 011; 174; 175; 176; 177; 178; 179; 180; 181; 182; 183; 184; 308; 309; 310; 311; 312; 313; 314; 315; 316; 317; 318; 319; 320; 321; 322; 323; 324; 325; 326; 327; 328; 352; 353; 354; 355; 356; 357; 358; 359; 360; 361; 362; 363; 461; 462; 463; 532; 588; 589; 590; 591; 592; 593; 594.

